# AntiDIF: Accurate and Diverse Antibody Specific Inverse Folding with Discrete Diffusion

**DOI:** 10.1101/2025.07.12.664553

**Authors:** Nikhil Branson, Charlotte Deane

## Abstract

Inverse folding is an important step in current computational antibody design. Recently deep learning methods have made impressive progress in improving the sequence recovery of antibodies given their 3D backbone structure. However, inverse folding is often a one-to-many problem, i.e. there are multiple sequences that fold into the same structure. Previous methods have not taken into account the diversity between the predicted sequences for a given structure. Here we create AntiDIF an **Anti**body-specific discrete **D**iffusion model for **I**nverse **F**olding. Compared with stateof-the-art methods we show that AntiDIF improves diversity between predictions while keeping high sequence recovery rates. Furthermore, forward folding of the generated sequences shows good agreement with the target 3D structure.

## 1. Introduction

Computational protein design has made significant progress with the development and utilisation of deep learning methods and generative AI e.g. (Geffner et al., 2025; Abramson et al., 2024; Cutting et al., 2024; Watson et al., 2023; Yim et al., 2023; Hsu et al., 2022). Antibodies are a particularly important type of protein. They are a large class of therapeutics and have been successfully used to improve human health for a host of different diseases e.g. (Lu et al., 2020; Carter & Lazar, 2018). Advances in machine learning have already and promise to continue to help in the development of novel antibody therapeutics (Jaszczyszyn et al., 2023; Hummer et al., 2022).

Inverse protein folding is the problem of finding amino acid sequences that fold into a given 3D protein backbone structure. Successful inverse folding models are invaluable in de novo design pipelines and for optimising desirable properties of proteins. Deep learning has proved highly effective at the inverse protein folding problem (Ingraham et al., 2019; Dauparas et al., 2022; Hsu et al., 2022).

Antibodies are Y-shaped proteins formed of two heavy and two light chains with the antigen binding site being mostly formed by complementary determining region (CDR) loops. These loops are hypervariable in comparison to the framework regions of antibodies that are mostly germline-encoded (Sela-Culang et al., 2013). The CDR loops are thus vital in determining the binding properties of antibodies. These distinct properties relative to other proteins mean that antibody-specific models have shown substantial improvement over general inverse folding models (Høie et al., 2023; Dreyer et al., 2023).

However, these models have focused on and have been evaluated for how well their predictions agree with the original sequences and by using forward folding methods, the consistency of the predicted structures with the target structures. Crucially this does not take into account the diversity between generated sequences, where because inverse folding is a many-to-one mapping problem, there are multiple sequences that can fold into a given structure. Having multiple diverse sequences is particularly valuable in practice as this gives a range of sequences that can be selected based on desired properties. For example, improved stability or reduced immunogenicity, Thus, it is desirable for an inverse folding method to generate a diverse set of plausible amino acid sequences for a given 3D structure (Silva et al., 2025; Yi et al., 2023).

Diffusion-based generative models have been shown to be particularly well-suited to producing a range of high-quality outputs in multiple domains including protein design (Dhariwal & Nichol, 2021; Song et al., 2020; Klarner et al., 2024; Yim et al., 2023; Watson et al., 2023). Thus, here we train an antibody-specific diffusion model for inverse folding (AntiDIF) to generate diverse and accurate sequences for a given backbone. AntiDIF is built upon RL-DIF (Ektefaie et al., 2024) a discrete diffusion inverse folding method that has shown good performance for general proteins. We train AntiDIF by fine-tuning RL-DIF with both experimentally determined structures from the structural Antibody Database (SAbDab) (Schneider et al., 2022; Dunbar et al., 2014) and predicted structures from the Observed Antibody Space (OAS) (Olsen et al., 2022; Kovaltsuk et al., 2018). We show that AntiDIF is able to generate diverse sequences while keeping high sequence recovery rates. Giving an unparalleled sequence recovery to diversity trade-off. Furthermore, we demonstrate that despite the improved sequence diversity forward folding of the sequences generated by AntiDIF maintains high levels of agreement with experimentally determined structures. These results show that AntiDIF would be a valuable addition to antibody design pipelines allowing for improved optimisation of biophysical properties.

## 2. Methods

### 2.1. Inverse folding with diffusion

Formally, the aim of inverse folding is to generate sequences *S* that fold into a protein backbone with the coordinates *X ∈* ℝ ^*N ×*3^, where *N* is the number of residues of the protein. Here, the amino acids are parametrised as onehot encoded vectors such that *S ∈{*0, 1*}* ^*N×v*^ with *v*, the vocabulary size, equal to 20, representing the 20 naturally occurring amino acids.

Diffusion consists of a forward noising process, where a denoising neural network can be learned, and a reverse process, used for generation. The distribution of noised states that relate the original native sequence *S*_0_ to latent noisy sequences is given by a Markov process such that:

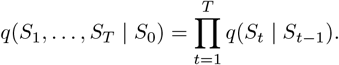

Where each latent state *S*_1_, *S*_2_, …, *S*_*T*_ is progressively noisier as the diffusion time step, *t*, increases, where *t ∈*ℕ_0_.

In the backward process, a noisy sample is iteratively denoised such that:

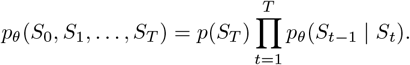

Here *p*_*θ*_ found via a neural network that is learnt to de-noise the latent states,and *θ* are the learnable parameters. The network steers the generation of new sequences toward the distribution of observed sequences.

We employ discrete diffusion to train AntiDIF as implemented in RL-DIF (Ektefaie et al., 2024) and based on D3PM (Austin et al., 2021). This defines a forward diffusion process by relating *S* at different diffusion time steps via transition matrices, which give the probability of moving from one amino acid type to another. See (Ektefaie et al., 2024) and (Austin et al., 2021) for full details.

The core layers of the de-noising neural network used in RL-DIF are PiGNN layers from PiFold (Gao et al., 2023). Given the backbone coordinates *X* the model creates a kNN graph with *k* = 30 between the residues. The PiFold featuriser is then used to obtain node and edge features that are used as input into the model with the noised sequence and the diffusion timestep. This model (*M*) thus, predicts the denoised sequence conditioned on the noisy sequence, backbone structure, and time step. Such that *Ŝ* = *M* (*S*_*t*_, *X, t*).

### 2.2. Training AntiDIF on antibodies

We utilised the dataset curated by Dreyer et al. for AbMPNN (Dreyer et al., 2023) for training and testing our model. Dreyer et al. utilised 3, 500 experimentally solved structures from SAbDab (Schneider et al., 2022) and 147, 919 (Olsen et al., 2022) predicted structures of sequences from OAS.

Clustering and filtering were applied to this data before splitting it into 80% training 10% validation and 10% testing sets. See Appendix A.1.1 for further details.

For each training epoch, we used all training samples from SAbDab and randomly sampled half this number of data points from the OAS training set to encourage diversity and limit overfitting. We trained until convergence on the validation set. We used a maximum learning rate of 1 *×* 10^*−*3^ with warm-up and decay. For our learning rate scheduler, we used a polynomial, of degree 2, and 1000 warm-up steps. We also used the adamW optimiser (Loshchilov & Hutter, 2019) and a batch size of 16. We started training from the pre-trained weights provided by RL-DIF (Ektefaie et al., 2024).

### 2.3. Inverse folding metrics

Previous evaluations of antibody-specific inverse folding methods have relied upon consistency metrics such as sequence recovery and structural agreement. However, as discussed in the introduction diversity between generated sequences is also a key metric. Therefore, in this study we report sequence recovery between generated and target sequences, diversity between generated sequences and RMSD between our generated structures, after forward folding, and the target structures.

For each backbone structure in the test set, we generate 4 sequences, to calculate the diversity, sequence recovery and RMSD from.

#### Sequence recovery

(SR) gives the fraction of amino acids in the generated sequence *Ŝ* that agree with the target sequence *S* for a given protein. Thus, for a protein with *N* residues, the sequence recovery is given by:

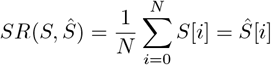

Where *S*[*i*] gives the amino acid identity at position *i* of the sequence.

#### Sequence Diversity

measures the fraction of amino acids that differ between a set of predicted sequences. For the set of *M* predicted sequences 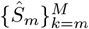 we calculate the diversity *D* between them as:

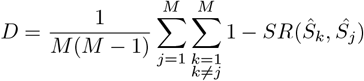

#### RMSD between 3D structures

After forward folding of our generated sequences, *F* (*Ŝ*) with forward folding method *F*, we find the root mean squared deviation (RMSD) between *F* (*Ŝ*) and the experimental backbone. We do this using the rsm cur function from PyMOL. We find the RMSD for the backbone atoms, after aligning the framework regions.

## 3. Results

### 3.1. AntiDIF shows high sequence recovery and diversity

Figure 1 shows the sequence recovery and diversity of AntiDIF compared with state-of-the-art inverse folding models (Høie et al., 2023). A table of these results is also provided in Appendix A.2. Figure 1A shows that AntiDIF improves upon RL-DIF, the general protein model we build on, for all CDRs. The figure also shows AntiDIF has a similar sequence recovery to AntiFold and has better sequence recovery than all of the other models across all CDRs, including the antibody specific model AbMPNN. Figure 1B shows that AntiDIF has substantially improved diversity of predicted sequences compared with AntiFold with a close to an order of magnitude improvement across multiple CDRs. Taken as a whole, these results demonstrate that AntiDIF greatly increases the diversity of predictions while keeping competitive sequence recovery rates.

**Figure 1.**
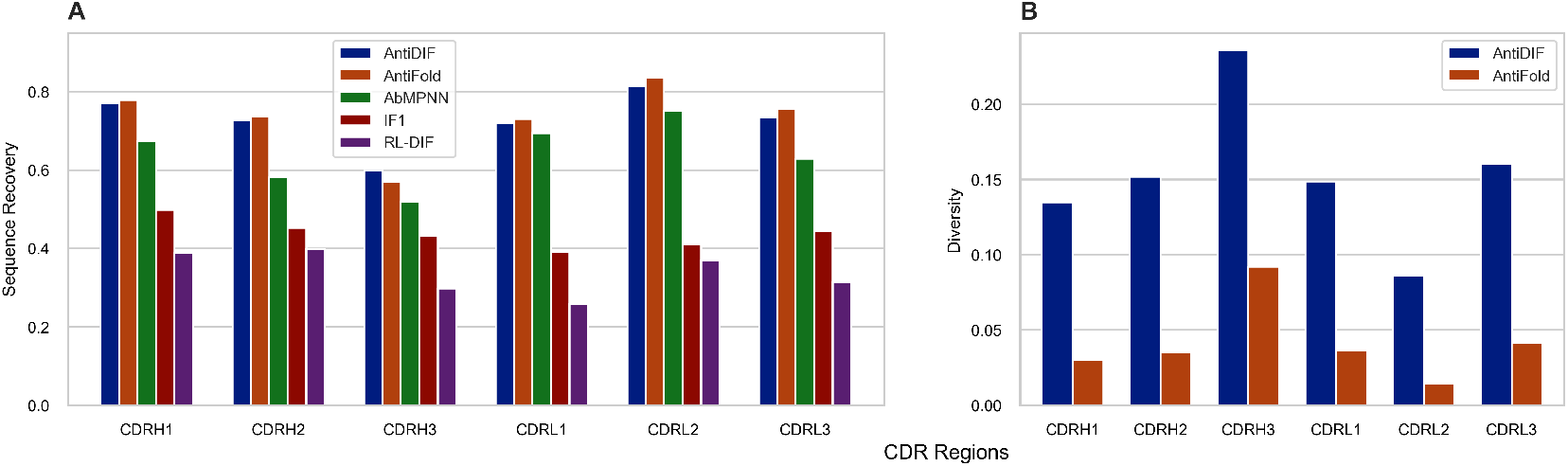
AntiDIF performance compared with antibody specific inverse folding methods, AntiFold and AbMPNN and general inverse folding methods ESM-1IF and RL-DIF. (A) Shows sequence recovery rates, for each CDR. (B) shows the diversity across four predicted sequences for the two models with the best sequence recovery, AntiDIF, and AntiFold. The figure shows AntiDIF has comparable sequence recovery to AntiFold while substantially improving diversity.

The methods achieve the lowest sequence recovery and the highest diversity for CDRH3. This is because CDRH3 is experimentally observed to be the most diverse loop due to junctional diversity (Weitzner et al., 2015).

Figure 2 compares AntiDIF with AntiFold sampled using higher temperatures than the default model (t=0.2). Figure 2A shows diversity against sequence recovery averaged over all CDRs. It shows, as expected, that increasing the temperature of AntiFold increases the diversity of the model’s predictions but this change also decreases the sequence recovery. Importantly, as temperature increases, AntiDIF outperforms AntiFold in terms of sequence recovery rates while still producing more diverse predictions. Thus, AntiDIF demonstrates a better trade-off between sequence recovery and diversity.

**Figure 2.**
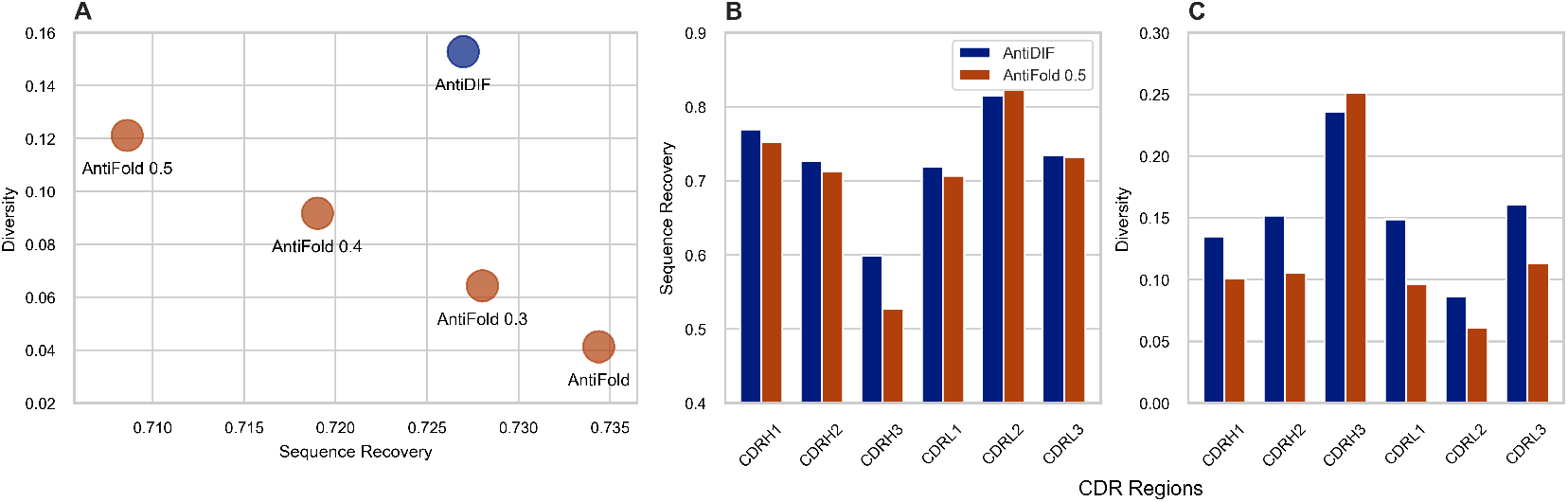
AntiDIF performance compared with increased AntiFold sampling temperatures (AntiFold default t=0.2). (A) shows diversity against sequence recovery averaged over all CDRs where the top right of the graph is where the best-performing model is located. (B) shows sequence recovery broken down by CDRs (C) shows diversity broken down by CDRs.

Figure 2B and Figure 2C break down the results by CDRs for AntiDIF compared with AntiFold 0.5. These figures show that AntiDIF has the highest sequence recovery across all CDRs apart from CDRL2 where it is still very competitive. Furthermore, AntiDIF has the best diversity across all CDRs apart from CDRH3, where it is slightly outperformed by AntiFold 0.5 which we note has greatly reduced sequence recovery for this region. AntiDIF also achieves both better sequence recovery and diversity across multiple regions showing a simultaneous improvement across both metrics of interest.

### 3.2. AntiDIF shows low RMSD of predicted 3D structures

Next, we evaluated how well our predicted structures compared with the experimental target structures from SAbDab. We did this by forward folding our predicted sequences using AbodyBuilder2 (Abanades et al., 2023). We then calculated the root-mean-square deviation (RMSD) for the backbone of the CDRs after aligning the structures on the framework region. For this analysis, we used the 56 structures with resolutions *<* 2.5 Å from the test set as was done in AntiFold for their structural agreement experiment (Høie et al., 2023). We considered the four predicted sequences per protein for each model.

Table 1 (and Figure 3 in Appendix A.2) shows these results across the different CDRs. The table shows that AntiDIF has a very similar performance to AntiFold across the majority of the CDRs. Furthermore, for CDRH3 AntiDIF has slightly lower RMSD despite AntiDIF having much higher diversity (Figure 1). Thus, AntiDIF generates both diverse sequences and plausible structures.

**Table 1.** RMSD (lower is better) between predicted structures from ABodyBuilder2 and experimentally determined backbones. Predicted sequences from AntiDIF and AntiFold are used for the respective models and the target (native) sequence is used for Native.

|  | CDRH1 | CDRH2 | CDRH3 | CDRL1 | CDRL2 | CDRL3 |
| --- | --- | --- | --- | --- | --- | --- |
| AntiDIF | 0.758 | 0.738 | 1.498 | 0.769 | 0.474 | 0.76 |
| AntiFold | 0.751 | 0.735 | 1.67 | 0.751 | 0.414 | 0.77 |
| Native | 0.69 | 0.585 | 0.846 | 0.577 | 0.355 | 0.568 |

**Table 2.** Sequence recovery of the different CDR regions (higher is better).

|  | CDRH1 | CDRH2 | CDRH3 | CDRL1 | CDRL2 | CDRL3 |
| --- | --- | --- | --- | --- | --- | --- |
| AntiDIF | 0.769 | 0.727 | 0.598 | 0.719 | 0.815 | 0.734 |
| AntiFold 0.5 | 0.752 | 0.712 | 0.527 | 0.706 | 0.822 | 0.731 |
| AntiFold 0.4 | 0.763 | 0.722 | 0.543 | 0.716 | 0.827 | 0.742 |
| AntiFold 0.3 | 0.771 | 0.731 | 0.559 | 0.725 | 0.833 | 0.75 |
| AntiFold | 0.778 | 0.736 | 0.57 | 0.73 | 0.835 | 0.756 |

**Table 3.** Diversity between the generated sequences for the different CDR regions (higher is better).

|  | CDRH1 | CDRH2 | CDRH3 | CDRL1 | CDRL2 | CDRL3 |
| --- | --- | --- | --- | --- | --- | --- |
| AntiDIF | 0.135 | 0.152 | 0.236 | 0.149 | 0.086 | 0.16 |
| AntiFold 0.5 | 0.101 | 0.105 | 0.251 | 0.096 | 0.061 | 0.113 |
| AntiFold 0.4 | 0.074 | 0.076 | 0.196 | 0.071 | 0.049 | 0.084 |
| AntiFold 0.3 | 0.052 | 0.056 | 0.143 | 0.049 | 0.026 | 0.061 |
| AntiFold | 0.03 | 0.035 | 0.092 | 0.036 | 0.014 | 0.041 |

**Figure 3.**
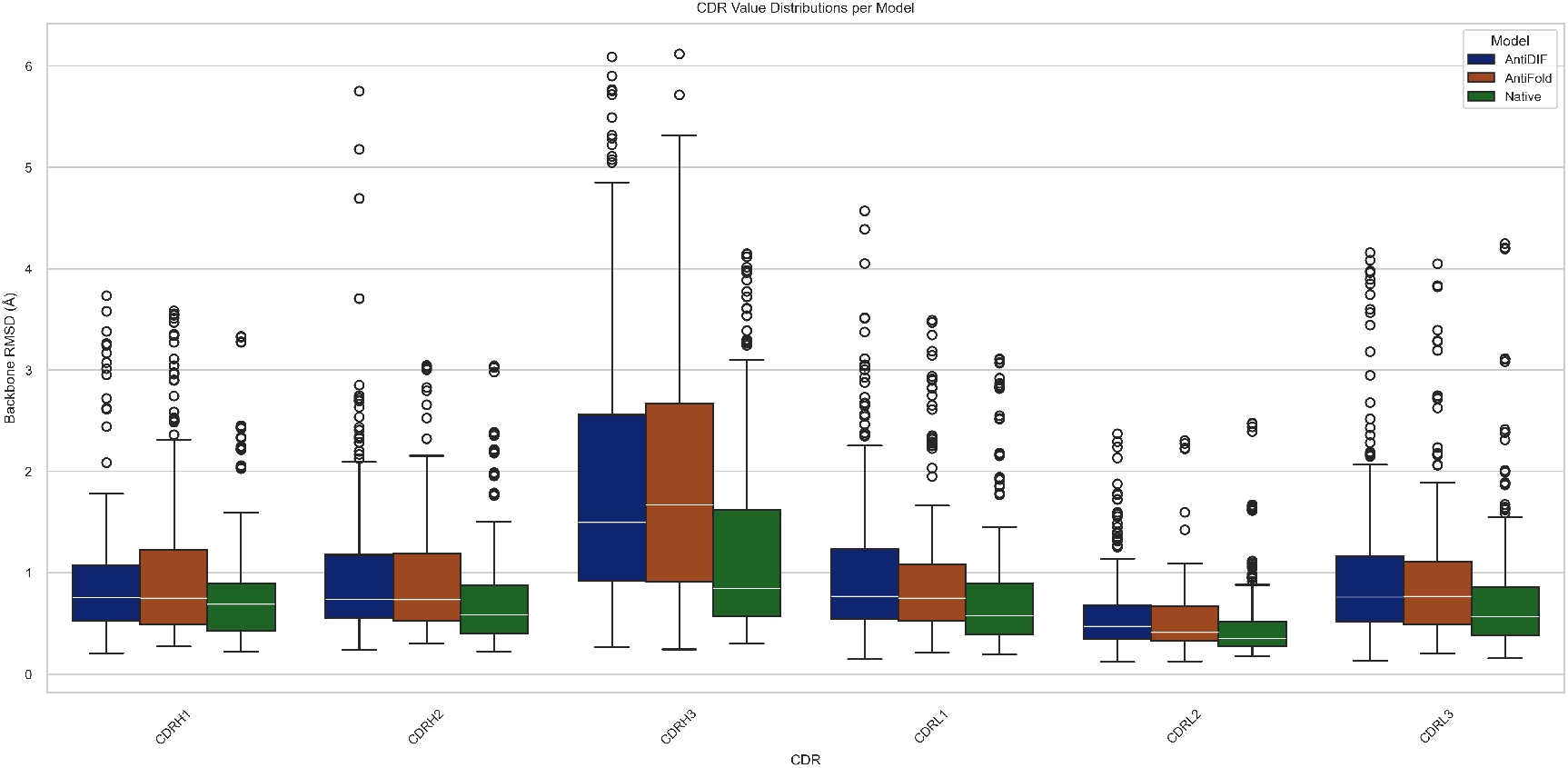
Box plot of RMSD (lower is better) across the 56 structures considered. RMSD is between predicted structures from ABodyBuilder2 and experimentally determined backbones.

## 4. Conclusions

Inverse folding is inherently a one-to-many mapping problem. Furthermore, having multiple sequences that fold into the same structure is particularly valuable in practice as this gives a range of sequences that can be selected based on important properties such as improved stability. However, diversity between predicted sequences has not been considered by previous antibody-specific methods. We demonstrate that by using discrete diffusion our model, AntiDIF, shows a better trade-off between sequence recovery and diversity. When, compared to state-of-the-art inverse folding methods, AntiDIF vastly improves diversity while keeping high levels of sequence recovery. Furthermore, the forward folding of our generated sequences shows good 3D structural agreement with the target structure. These results show that AntiDIF would be a valuable addition to antibody design pipelines.

## Software and Data

The code is available from https://github.com/oxpig/AntiDIF. The data used is all publicly available (Dreyer et al., 2023).

## Acknowledgments

Thank you to Oliver Turnbull, Alissa Hummer, Frédéric Dreyer, Magnus Høie and Yasha Ektefaie for their helpful feedback and discussions.

for their helpful comments and discussions.

## Impact Statement

This paper aims to advance the field of Machine Learning for protein design. We hope it can be used to help improve the discovery process for novel antibody therapeutics to help improve healthcare.

## A. Appendix

### A.1. Additional methods

Here we briefly detail discrete diffusion for inverse folding using D3PM see (Austin et al., 2021) and (Ektefaie et al., 2024) for full details.

The forward noising process is defined for amino acid sequence *S* at time step *t*, such that

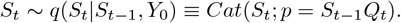

Here *Q*_*t*_ is a transition matrix, of dimensionality *v × v* and *S*_0_ is the original noise-free sequence and *Cat* is the categorical distribution. As in RL-DIF we use uniform transition matrixes.

To parameterise the reverse process for generation we also need the posterior given by:

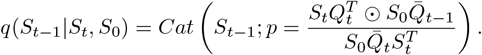

where 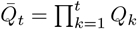. Thus, using a neural network allows for the prediction of the sequence *S*_0_ and thus, *q*(*S*_*t−*1_|*S*_*t*_, *S*_0_).

Therefore, after sampling *S*_*T*_ from a noised distribution, *S*_*T*_ can then be iteratively de-noised to produce a clean sample *S*.

#### A.1.1. DATASETS

The SAbDab structures selected by Dreyer et al. are antibodies in complex with antigens. As in AntiFold we only used the antibody fragments in this study. These antibodies are numbered using IMGT numbering (Lefranc et al., 2003). The OAS dataset does not contain epitopes. Dreyer et al. filtered the SAbDab structures to remove redundant structures and ones with high levels of CDR or epitope sequence similarities. They also only keep structures with experimental resolutions *<* 5Å. The OAS dataset was filtered by removing duplicates. These datasets were then clustered based on the similarity of the CDR sequences. The train test split was done based on these clusters such that a SAbDab cluster that would have been in the OAS training or validation clusters was in the SAbDab training set to ensure no overlap. We further filtered this dataset by removing structures that could not be modelled by RL-DIF. This lead to the removal of 11 data points from the test set and 32 from the train set. The results for all models presented here are from this same test set.

### A.2. Additional figures and tabels

## Notes

### Competing Interest Statement

The authors have declared no competing interest.

### Funding

- Wellcome Trust

